# Optogenetic activation of entorhinal projection neurons alters the target recognition and circuit development without enhancing axon regeneration after axotomy in organotypic slices

**DOI:** 10.1101/2025.06.22.660916

**Authors:** Karen Wells-Cembrano, Marc Riu-Villanueva, Juan José López-Jiménez, Miriam Segura-Feliu, Rosalina Gavín, José A. del Río

## Abstract

The central nervous system (CNS) has a limited intrinsic capacity for axonal regeneration, making functional recovery after injury extremely challenging. Numerous strategies have been explored to overcome this blockade, among others, molecular interventions or modulation of the inhibitory extracellular environment. Despite some advances, effective regeneration remains elusive, particularly in adult CNS neurons. To investigate these mechanisms in a controlled and reproducible setting, we employ organotypic slice cultures (OSCs), which retain key structural and cellular features of the intact brain while allowing for long-term *in vitro* experimentation. In particular, the entorhino-hippocampal (EH) co-culture model preserves the anatomical and functional connectivity of the perforant pathway, providing an excellent platform for studying axonal degeneration and regeneration. This model reproduces laminar specificity, axonal myelination, and inhibitory signaling after axotomy, closely mimicking *in vivo* conditions. Furthermore, EH co-cultures facilitate the application of optogenetic tools to monitor and manipulate neuronal activity. Our study explores whether enhancing activity in entorhinal cortex neurons can promote axonal regeneration after a EH lesion. Our results show that increased activity in entorhinal neurons alters the development of the EH connection and fails to enhance the regrowth of injured mature entorhinal axons. These findings suggest that both extrinsic and intrinsic factors shape the regenerative response and highlight the utility of EH OSCs as a versatile model for testing future pro-regenerative interventions.

## Introduction

Perinatal brain slice cultures offer unique advantages over other 2D or 3D *in vitro* methods, since they mimic numerous *in vivo* aspects due to the presence of most of the cell types observed *in vivo* [1–3]. For most purposes, slices obtained from early postnatal brain and cultured for long period (>20 days), termed organotypic slice cultures (OSC), largely preserve a high degree of cellular differentiation and tissue organization (e.g., laminar organization and synaptic target specificity) during culture (see below). OSCs have been prepared from several brain regions, including hippocampus, neocortex, allocortex. striatum, spinal cord, hypothalamus or cerebellum [4–15]. Relevantly the use of OSCs obviate the need for extensive animal surgery and expensive laboratory equipment (e.g., [16]). Thus, their use in basic and applied research has been develop over the years and more recently due to the 3Rs strategy of animal research [1, 14, 15, 17]. In fact, OECs showed intrinsic relevance that also shared other approaches based in the use of induced pluripotent stem cells (iPSC) such us 3D brain organoids [18–20]. As advantages, a number of neuropathological events (from genetic, traumatic or infective) that affect specific brain regions have been easily reproduced in OECs [21–32] being currently also used as model of neurodegeneration (e.g., Alzheimeŕs [33–35] or Parkinsońs [36] diseases). Several methods and modifications have been developed for long-term culture of brain slices, from the pioneer *roller tube* technique of Gahwiler [37] to the interphase method of Muller and coworkers [1, 11, 38] or current methods using adult-derived brain tissue (e.g., [39–41]. Pioneer OSCs [37] have proved tedious to prepare with high cytoarchitectonic variability due to thinning of slices. Furthermore, hydrogel-embedded cultures show intense glial reactivity, which largely conditioned functional studies and optical *in situ* analysis [38, 42]. The membrane interface culture method (MICM) developed by Muller and coworkers facilitates access to the culture (from optical to pharmacological studies) (see [1, 11, 16] for reviews). This culture technique and its current modification allow the culture nerve cells (from neurons to glial cells) that are highly differentiated in terms of their morphological and physiological characteristics (see [5, 43, 44] for classic studies). Moreover, the presence of glial cells is believed to provide a microenvironment that facilitates differentiation of neurons. In this respect, glial (mainly astroglial) cells proliferate in the culture and a border of laterally migrating reactive astrocytes surrounds the MCIM [38, 42, 43]. Scarred astroglia cells in MICM cultures are also located at the bottom of the slice. This is important, since the culture medium is applied below the membrane and this dense glial scar hinders the putative drug treatment and, in some studies, additional functional analysis [45, 46].

In anatomical terms, rodent hippocampus and the dentate gyrus are discrete brain regions with a unique laminar organization of cell layers and afferent connections [47]. The entorhino-hippocampal connection (EHC), also termed the perforant pathway, is the main afferent connection to the hippocampus. The perforant pathway derives from neuronal cells located in the upper layers of the entorhinal cortex (mainly layer II-III), which terminate in the stratum lacunosum-moleculare of the hippocampus proper and the outer two-thirds of the molecular layers of the dentate gyrus [48, 49]. The EHC has been analyzed in detail and various factors (cellular and molecular) mediating its development have been determined (see [50] for review). Moreover, as seen in pioneer studies by G. Raisman [51], P.L. Woodhams [52, 53] and M. Frotscher [54, 55], reviewed in [12, 15] the entorhino-hippocampal (EH) co-cultures have some key and useful features that make them particularly interesting in studies of axonal regeneration: i) the EHC is reproduced easily *in vitro* in MCIM cultures (see above references); ii) the EHC *in vitro* showed a laminar specificity similar to that found *in vivo* (e.g., [12, 50, 56–58]); iii) the EHC is myelinated both *in vitro* in MCIM similarly as *in vivo* (e.g. [59–61]); and iv) most of the cellular and molecular barriers to axon regeneration (e.g, myelin or chondroitin sulphate proteoglycans) are present or overexpressed by different cell types after the axotomy of the mature perforant pathway *in vitro* similarly as *in vivo* (e.g., [38, 59, 62, 63]). Thus, due to the target-specific termination of the EC axons, it is possible to study their growth or remodelling and to quantify the number of reinnervating axons after axotomy *in vitro* (e.g., [58]). Indeed, this platform has been used to demonstrate age-related decline in the ability of EH axons to regenerate, as occurs *in vivo* during postnatal development [42, 52, 53, 61, 63–65]. The ability of young hippocampal tissue to support the regrowth of lesioned E18-P1 entorhinal axons in culture has been examined in detail for many years by using heterochonic co-cultures (e.g., [42, 52, 53, 64, 65]). This young hippocampus appears permissive for regeneration and results in an axonal projection to the hippocampal tissue terminating in the appropriate targets [42, 52, 53, 64, 65]. However, it has also been established that there is a limited ability to modify the regenerative connections made from the postnatal entorhinal cortex by juxtaposition of different hippocampal target fields due to the apparent early specification of entorhinal projection neurons [66–68]. In addition, most of these cues disappeared progressively from the hippocampal tissue during the first 10-15 days in culture [60, 65]. Thus, because of this limited capacity for regeneration following lesion within the perforant pathway the period at 10-15 DIV provides an optimal basis for screening agents and modifications that might enhance axon regeneration. On the other hand, not only the inhibitory environment plays a role in the absence of regeneration in OECs. In OECs, the projecting EC neurons survive after axotomy at 15-21 DIV but are unable to regrowth [65, 69]. Thus, in the last years there have been several studies approaching different aspects of the projecting neurons physiology aimed to trigger the intrinsic regrowth capacity [70, 71]. In this direction, the role of the neuronal activity modulation has been explored. Thus, for spinal cord, electrical stimulation has been shown to enhance regeneration of sensory axons after peripheral nerve or dorsal columns injury; and sprouting of cortical axons into contralateral spinal cord grey matter after axotomy (e.g., see [72] for details).

Taking all this into account and the properties of the OECs, in this study, we will focus our attention in OSC containing EC and hippocampus to explore axonal development and regeneration of CNS neurons after axotomy and whether this regeneration can be modulated by neuronal activity using projecting EC neurons as model. Using cell-directed optogenetics, calcium imaging and axonal tracing techniques, axonal development and regeneration will monitored in MCIM cultures were some of the molecular and cellular mechanisms responsible for its absence after adult CNS lesions are described [59, 65, 69, 73, 74].

## Material and Methods

### EHC experiments

Cocultures were prepared from P0-P1 CD1 newborn mice (Charles River, Lyon, France). After brain removal, single horizontal sections (300-350 μm thick) containing both the EC and the hippocampus were obtained using a tissue chopper (Mickle Laboratory Engineering, Gomshall, UK) and maintained in ice cold slice preparation media (75% minimal essential medium (MEM), 25% HBSS, 2 mM glutamine and 0.044% NaHCO3, pH 7.3). Slices were further dissected under dark-field binocular control to isolate the hippocampus and the EC. In dissected slices the subiculum was preserved with the hippocampal slice. After dissection, 50% of the dissected EC (n= 36) were incubated overnight under gentle agitation with AAV9-Syn-ChR2 (pAAV9.Syn.GCaMP6s.WPRE.SV40, Addgene cat. 100843-AAV9, [75]), virus (diluted 1:500 v/v) and the remaining ECs were mock transfected. In some cases. Some EC were simultaneously double infected with AA9-Syn-ChR2 and AAV9-Syn- JRCaMP1b (AAV9.Syn.NES-jRCaMP1b.WPRE.SV40, Addgene cat. AV-1-PV3852, [76]).

The dissected hippocampal slices were seeded onto Millicell^TM^ transwells (PICM0RG50, 5 mmm high and 30 mm diameter, Merck Life Science S.L.U., Madrid, Spain) and incubated with culture media (see below). Next day, the infected and non-infected ECs were rinsed in incubation media and the EHC were stablished by transferring the EC slices to the cultured hippocampus and reoriented using fine tungsten needles. Cocultures were fed with 1.2 ml of culture medium (50% MEM, 25% horse serum, 25% HBSS) containing 2 mM glutamine and 0.044% NaHCO3, pH 7.3. The membrane cultures were maintained in a humidified incubator at 37°C in 5% CO2. This day was considered 1 DIV. The medium was changed after 24 h and subsequently every 48 h until the EHC was examined. Two groups of EHC cultures were stablished for experiments. A group of 18 EHC were stimulated with blue (470 nm) light (n = 9 AAV9-Syn-ChR2 infected and n = 9 uninfected; see below). In a second group the EH connection was axotomized at 7 DIV (n= 9 AAV9-Syn-ChR2 infected and n = 9; uninfected) and 14 DIV (n= 9 AAV9-ChR2 infected and n = 9; uninfected) by cutting the cocultures from the rhinal fissure to the ventricular side along the entire EH interface with a thin tungsten needle [65]. This second group was also stimulated with blue light following the protocol described below. The first group was processed after 7 DIV (5 days of light treatment). The second group (axotomized at 14 DIV) was processed after 7 post-light treatment. All experiments were performed under the guidelines and protocols of the Ethical Committee for Animal Experimentation (CEEA) of the University of Barcelona, and the protocol for the use of animals in this study was reviewed and approved by the CEEA of the University of Barcelona (CEEA approval #276/16 and #141/15). Methods are reported in accordance with the Animal Research: Reporting of In Vivo Experiments (ARRIVE) guidelines.

### *In vitro* optogenetic stimulation of EC projecting neurons

EHC containing the infected ECs with AVV9-Syn-ChR2 were used for *in vitro* optogenetic stimulation experiments. A 470 nm emission LED array (LuxeonRebel^TM^) under the control of a Driver LED (FemtoBuck, SparkFun) of 600mA and a pulse generator PulsePlus (Prizmatix, Israel) was used to deliver blue light to neuronal cultures [77]. During experiments, temperature changes inside the optogenetic platform were monitored by an Arduino-UNO™ microcontroller using a temperature probe DS18B20 with PWM relay output (CEBEK I-86, Spain) connected to a 12 V cooling fan. The cooling fan switched on with temperature increases >0.5 °C from 37 °C. The LED array was placed onto aluminum heat sink plates below the culture dishes in the optogenetic platform at a distance of 2 cm to ensure that the complete area of the 35 mm well plate was illuminated. We avoided to use long prolonged stimulations protocols [78] to ensure cell survival after stimulation. The optogenetic stimulation protocol consisted in cycles 1 h of illumination at 20Hz of frequency with 5-45 ms pulses, in 1s ON-1s OFF periods [72]. In these conditions, the external stimulation voltage unit drives 12 V and 600 mA for each Quad LED module, with an average light intensity of approximately 20-25 mW/cm^2^ measured at the culture dish containing the devices with a Newport 1919 optical power meter (Newport Photonics, USA). The stimulation was applied 24 h after the establishment of the EHCs (see above). For axotomized EHC slices the stimulation time-point after the axotomy was 24 h after axotomy and developed twice a day (12 h interval) during 7 days as above indicated.

### Calcium imaging in AAV9-Syn-ChR2 and AAV9-Syn-jRCaMP1b double infected EC neurons

Some double-infected (AAV9-Syn-ChR2 + AAV9-Syn-jRCaMP1b) EH cocultures were analyzed for colocalization and Ca^2+^ analysis in the EC in an Olympus IX61 microscope equipped with an Fluoview-II-CCD monochrome (1.42 effective MPx) or a DP71-CCD RGB (1.5 effective MPx) cameras and Cell^P^ software (Olympus; Japan). The microscope used a X-Cite 120 PC Q illumination system (Xenon halide lamp of 100W working at a 50% regime, Excelitas Noblelight, Alzenau, Germany). Double immunofluorescence images were obtained to demonstrate that stimulated EC neurons expressed both, ChR2 and jRCaMP1b. The analysis of the colocalization in both 8 different coinfected cultures was developed analyzing paired fluorescence images (1024 x 1024 x pixels, DP71 camera) at the green and red channels using the “Just Another Colocalization Plugin” (JACoP) Fiji^TM^ plugin [79]. For optogenetic stimulation, slices were illuminated at 15 DIV using *ad hoc* designed filter sets by using a 20X water immersion objective (Olympus UMPlanFL, NA= 0.5). Pictures (512 x 512 pixels, Fluoview II camera) were obtained each 200 ms, allowing to capture Ca^2+^ changes revealed by jRCaMP1b. Thus, in the experiments jRCaMP1b basal fluorescence changes was recorded without optical stimulation during 5-10 sec. After this time, the slice was illuminated for 30 sec with a 470 nm diode laser collimated beam of 0,4 mm diameter and 25 mW at 90° with respect optical axis. The xyz position of the laser bean was controlled using a 3-axis micromanipulator (Nashirigue, Japan) to the EC and the changes induced by the illumination recorded for additional 40 sec (80 sec per video in total). Time lapse videos (multi *.tif files) were mounted in Fiji^TM^, and denoised using Aydin^TM^ v.0-1-13 https://doi.org/10.5281/zenodo.5654826, with the self-supervised auto-tuned Classic-Butterworth algorithm, and motion corrected and ROIs selected by using EzCalcium [80] and NeuroSeg[81]. Thus, the average fluorescence *Fi (t)* in each ROI *i* along the recording was then extracted, corrected for global drifts and artifacts, and finally normalized as *(Fi (t) — F (0,i)) / F (0,i) = fi (t)*, where *F 0,i* is the background fluorescence of the ROI. The time series of *fi (t)* was analyzed to determine sharp calcium transients. To reveal basic data on neuronal activity, normalized data files were first processed using the protocols and algorithms developed by Sun and Südhof [82] in Matlab^TM^ R2022b (lic. 48811179) [82]. Computation was developed in an iMac computer (macOS Big Sur 11.6, 3.2 GHz i7, 6 nuclei and 32 GB) running Python 3.9-12, Conda 23.5.2; Napari 0.4.12 and Matlab 2022b update 3 (9.13.0.2126072).

### EHC labelling with Biocytin

After treatments, a small crystal of Biocytin (Sigma, Poole Dorset, UK) was injected in the entorhinal slice to label the EHP. The following day, cocultures were fixed with 4% buffered paraformaldehyde for 4 hours and 50-μm-thick sections were obtained using a vibratome (Leica VT1000, Barcelona, Spain). Free floating sections were blocked using 5% fetal bovine serum diluted in 0.1M PBS containing 0.4% Triton X-100, rinsed in 0.1 PBS incubated overnight with an Avidin-Biotin peroxidase Complex (ABC-elite™, Vector Laboratories, Burlingame, CA, USA, diluted 1:100 in 0.1 M PBS, 5% fetal bovine serum and 0.5% Triton X-100;) and peroxidase activity was visualized using a nickel-enhanced diaminobenzidine (DAB) reaction. After DAB development, some cocultures were osmified and processed for electron microscopy. In addition, some cultures were double processed for Biocytin detection and Calretinin (a marker of hippocampal Cajal-Retzius cells) as indicated [65]. Only cultures displaying equivalent Biocytin labeling in the entorhinal cortex were considered in the study.

### Immunohistochemical procedures

For immunocytochemistry, vibratome sections (50 μm thick) were permeabilized with 0.5% phosphate buffered DMSO and blocked with normal serum containing Fab IgG fragments (Jackson Immunocytochemical Laboratories, West Grove, PA). Free-floating sections were incubated for two days with either anti-GFAP (diluted 1:2000), anti-NG2 (diluted 1:1000, Chemicon, International, Temecula, CA), or anti-MAG2 (diluted 1:200, Sigma) primary antibodies and the ABC Kit (Vector Laboratories). Controls, including omission of the primary antibody or its substitution by normal serum, prevented immunostaining. In some cases, EH cocultures infected with AAV-Syn-ChR2, were fixed with 2% buffered paraformaldehyde for 2 hours rinsed in 0.1M PBS and mounted in Mowiol^TM^.

### EHC treatments

Taking advantage of the early generation of Cajal-Retzius cells with respect to granule cells and pyramidal neurons and of their layer-specific distribution, we ablated Cajal-Retzius cells by local application of domoic acid (a-amino-3-hydroxy-5-methylisoxazole-4- propionic/kainate (AMP/KA) receptor agonist [57]. After two days the 95% of the Cajal-Retzius cells were removed from the hippocampal slice that received the EC slice. [57]. In a second of experiments, EHC were prepared and axotomized at 10 DIV. After extensive rinsing two small pieces of newborn (P0) hippocampus containing the *stratum lacunosum moleculare* and the molecular layer of the dentate gyrus *(slm/ml)* were transplanted in the lesion site [65]. After 7 additional DIV, cultures were traced with Biocytin and double processed for Biocytin detection and Calretinin staining as above.

### Quantitative Real-Time PCR on axotomized EC

The lesioned EC (stimulated *vs* non-stimulated, n=4 each) were harvested from the membranes by a fine spatula. Total RNA was then extracted using a RNeasy kit (Qiagen, USA), according to the manufacturer’s guidelines. cDNA was then synthesized with SuperScript™ II reverse transcriptase (ThermoFisher Scientific) from 1 mg of RNA per sample. Real-time qPCR was run with Light cycler 480 SYBR Green Master (Roche, Spain) in a StepONEPlus light cycler (ThermoFisher Scientific). Cts were calculated following the manufacturer’s instructions. Expression values are expressed as 2^-ΔΔCt^. First Cts were normalized versus glyceraldehyde-3- phosphate dehydrogenase (*GAPDH*) as a housekeeping gene. Primer sequences used are the following:

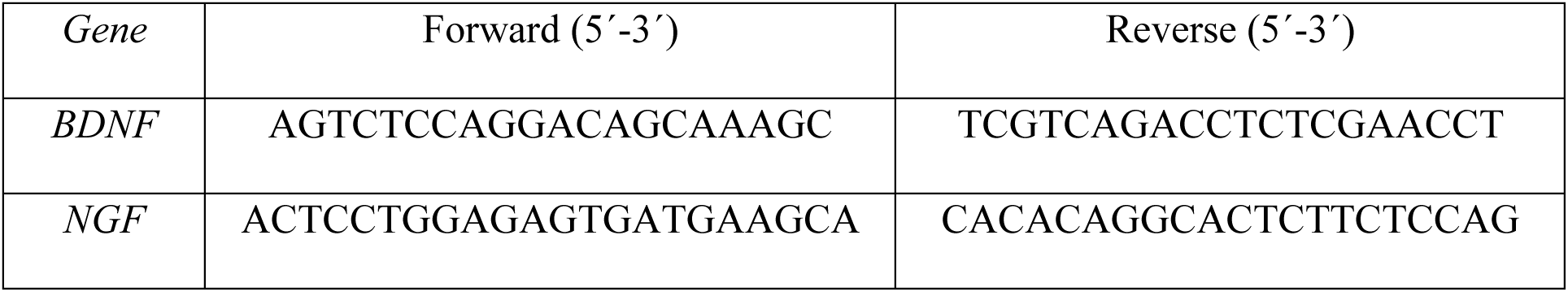
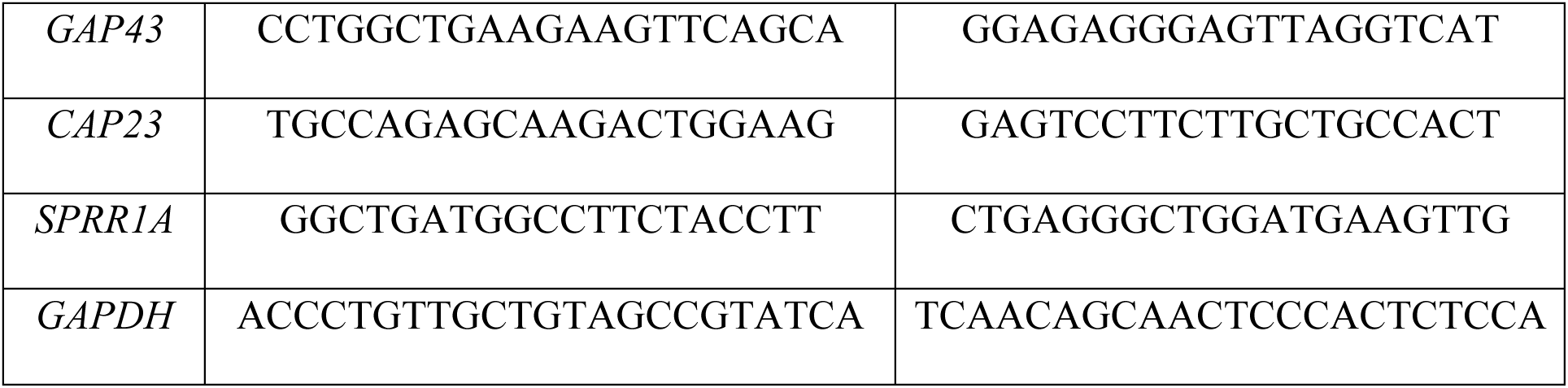

## Results

### Analysis of the regenerative properties of the lesioned EHP in OECs

OECs of the EHCs were prepared as indicated (Figure 1A). After 7-10 DIV Biocytin traced EHC displayed EC axons crossing the subicular region and innervating in a lamina specific manner the *stratum lacunosum moleculare (slm)* of the hippocampus proper and the molecular layer (*ml*) of the dentate gyrus (Figure 1B). Indeed, double labelled cultures with Biocytin and Calretinin showed that EC axons specifically innervate regions populated with hippocampal Cajal-Retzius cells (Figure 1C-D) as reported in other studies (e.g. [57, 83, 84]). Next in a second set of experiments, EHC were axotomized at 7 or 14 DIV, 12 days after axotomy (DAL) were traced with Biocytin and processed to light or electron microscope (Figure 1E-H). Revealed sections demonstrated the decline in the number of EHC that showed EC axon regeneration in the hippocampus and the dentate gyrus when axotomy was performed at 14 DIV (Figure 1E-F). In fact, none of the EHC lesioned at 14 DIV showed EC axons in the hippocampus (Figure 1F). In contrast, EHC lesioned at 7 DIV showed regenerating EC axons in the natural target regions. At electron microscopy, Biocytin labelled axon terminals were observed to contact with dendrites of pyramidal and granule cells in the lesioned group at 7 DIV (Figure 1G-H). In conclusion, our results demonstrate that layer specific regeneration of lesioned EC neurons is absent after 2 weeks *in vitro*.

**Figure 1.**
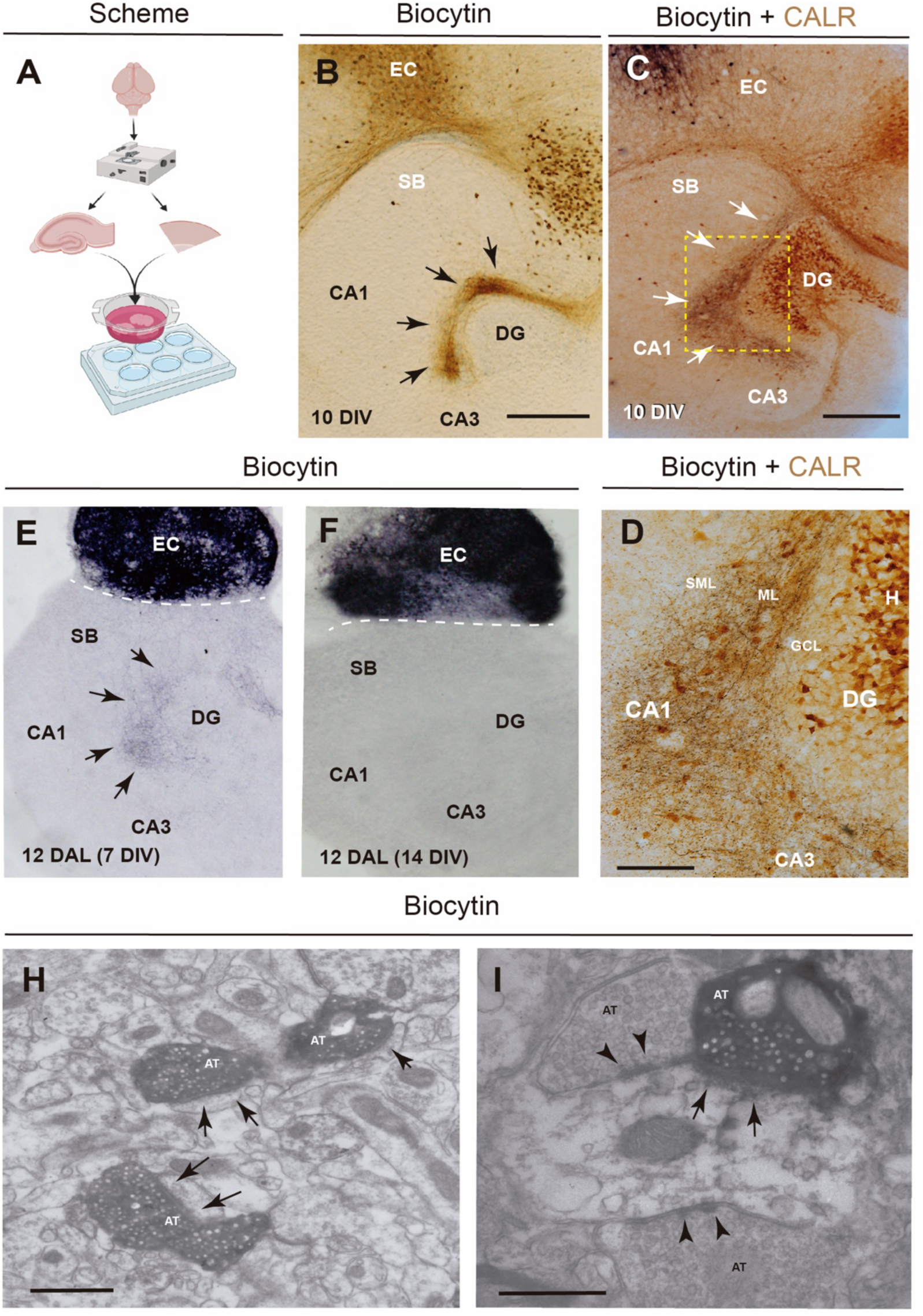
Decline of axon regeneration of EC axons in aged EHC. **A)** Scheme illustrating the procedure of the EHC preparation (see Material and Methods for details) **B)** Vibratome section of an EHC of 7 DIV labelled with Biocytin in the EC. Labelled Axons (arrows) innervate the hippocampus in a layer specific manner. **C-D)** Vibratome section of a double labeled EHC (Biocytin (black) and Calretinin (brown) of 10 DIV. The boxed region in **C** can be seen in **D**. **E-F)** Low magnification of Biocytin traced EHC 12 days after axotomy (DAL) at 7 **E** and 14 DIV **F**. Notice the regeneration of the EHC after axotomy at 7 DIV (arrows in E) and the absence of regeneration after a lesion at 14 DIV. **H-I)** Examples of electron microscopy photomicrographs illustrating the presence of Biocytin labelled axon terminal in the slm/ml of EHC lesioned at 7 DIV and analyzed 12 DAL. Arrows point to asymmetric synapses on unidentified dendrites. Arrowheads in **I** point to a symmetric contacts onto a dendrite that also showed a symmetric contact from a regenerating EC axon labelled with Biocytin. Scale bars: B = 200 μm pertains to E-F; C = 200 μm; D = 100 μm. H-I = 1 μm. Abbreviations: DG: dentate gyrus; CA1-2: Hippocampal regions; SB: Subiculum; EC: Entorhinal cortex; GCL: granule cell layer; ML: molecular layer; SLM. *stratum lacunosum moleculare*; H: Hilus; AT: Axon terminal.

### Increased glial scar generation after EHC lesion *in vitro*

The EHC axotomy *in vitro* was followed by a robust glial reaction as observed by immunocytochemistry (Figure 2). Slight cavitation (∼25-30 μm) occurred during the first 5-7 DAL, with abundant reactive astroglia (GFAP-positive; Figure 2B and D), ameboid microglia (not shown) and NG2-positive oligodendrocyte progenitors on both sides of the transection (Figure 2C and E). After 10 DAL, the cavity was filled with a dense network of glial cells and processes (see Figure 2E). Over-expression of the myelin associated glycoprotein (MAG) in small size proliferating oligodendrocytes after lesion was also observed in lesioned OECs at 15 DIV.

**Figure 2.**
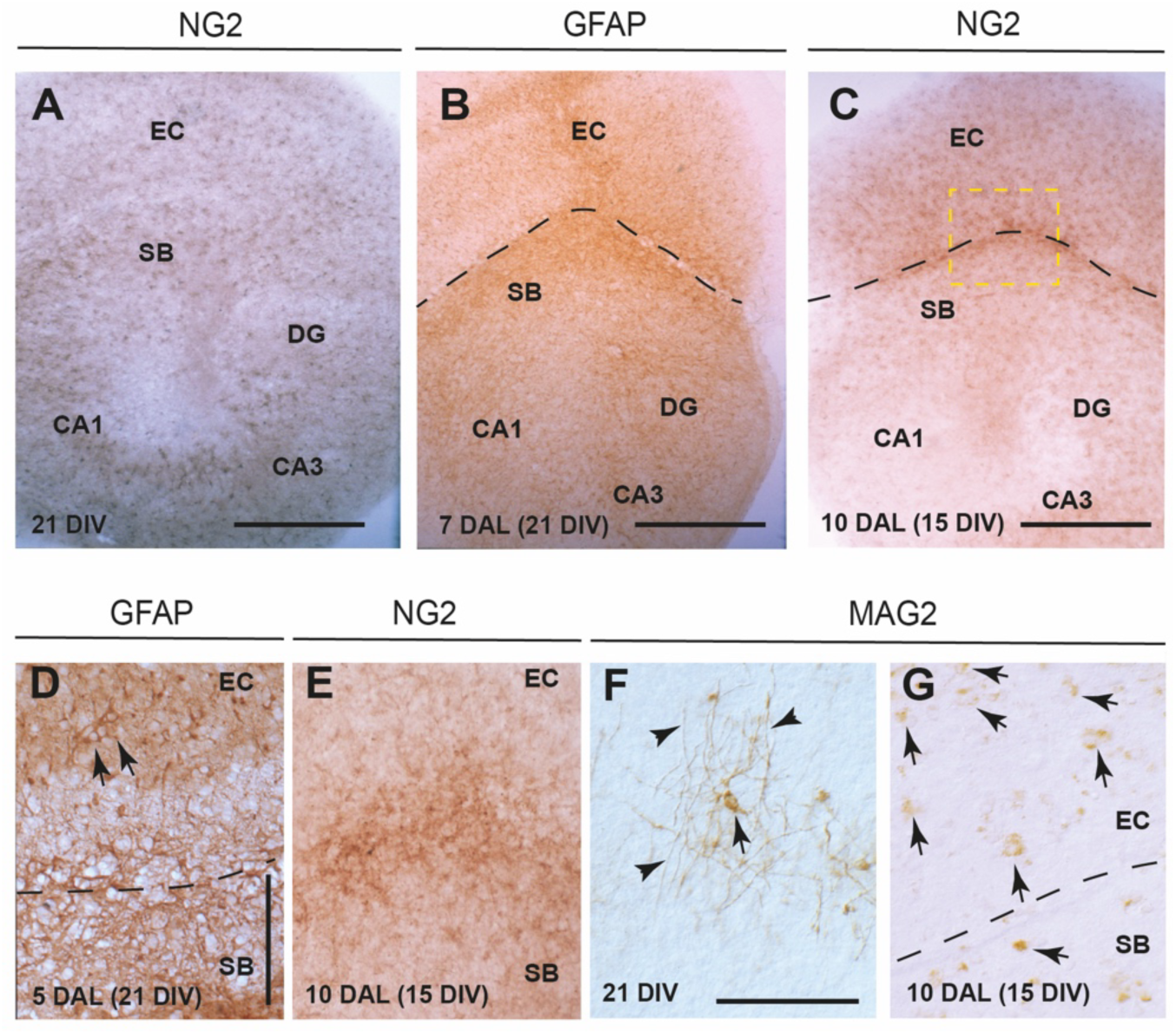
Glial proliferation after EHC axotomy *in vitro*. **A)** Low power photomicrograph illustrating the distribution of the NG2-positive astrocytes in a OEC after 21 DIV. **B-C)** Changes in GFAP (B) and NG2-immunostaining (C) after EHC axotomy at 21 DIV **B** and 15 DIV **C.** The dashed line showed the axotomy trajectory, and the boxed area in **C** is shown in **E**. Numerous reactive cells are located close to the axotomy transection (arrows in D). As indicated, a cavitation can be seen during the first week after lesion **D**, that further is occupied by reactive cells **E**. **F-G)** Example of the morphological changes in the MAG-positive cells after EHC axotomy. In non-lesioned cultures MAG positive cells (arrow in **F**) are mainly located in the white matter displaying long expansions (arrowheads in F) as chandelier-like processes. After axotomy **G,** these features are no longer observed, and a numerous number of small proliferating cells arrows with MAG-positive labelling in the soma can be observed in both sides of the lesion. Scale bars: A-C: 200 μm; D and F: 100 μm pertains to E and G respectively. Abbreviations as in Figure 1.

Target ablation of EC axons by domoic acid impair axon development of the EHP, but young target transplantation is unable to overcome the absence of EHP regeneration after axotomy in aged OECs.

Next, we aimed to explore whether changes in the target regions of the hippocampus could affect the regenerative potential of lesioned EC. Thus, as control we treated hippocampal slices with domoic acid to reduce the number of Cajal-Retzius cells in the hippocampus following the protocols published in [57, 85] (Figure 3A). 10 days after chemical lesion (DAL) the EHC was traced with Biocytin and the EHC were double processed for Calretinin and Biocytin (Figure 3B). Results demonstrated that the absence of the Cajal-Retzius cells impair the establishment of the EHC. In contrast, some subicular cells were able to innervate the entorhinal cortex being retrogradely labeled with Biocytin (Figure 3B). This situation was similar to those observed in OECs axotomized at 15 DIV al analyzed 10 DAL (Figure 3C). Next, we aimed to explore whether the ectopic transplantation of the postnatal *slm/ml* was enough robust to recapitulate the EC axon regrowth after lesion (Figure 3D). Thus, EHC lesioned at 10 DIV were transplanted with P0-dissected *slm/ml* pieces (Figure 3D-F). 7 DAL, OEC were processed for Biocytin labeling and Calretinin immunostaining. Results showed the survival of Cajal-Retzius cells in the transplanted *slm/ml* (Figure 3E-F). However, although some small bundles of EC axons were observed in the boundary, most lesioned EC axons were unable to enter in the transplanted target region as well in the remaining hippocampus. In contrast, some Cajal-Retzius cells (arrow in Figure 3F) were retrogradely labeled with Biocytin suggesting their capability to innervate the lesioned EC as suggested during development [86, 87]. This suggests that the presence of the appropriate target is not sufficiently robust to induce the regeneration of lesioned EC axons, and that other intrinsic factors might impair the expected regeneration.

**Figure 3.**
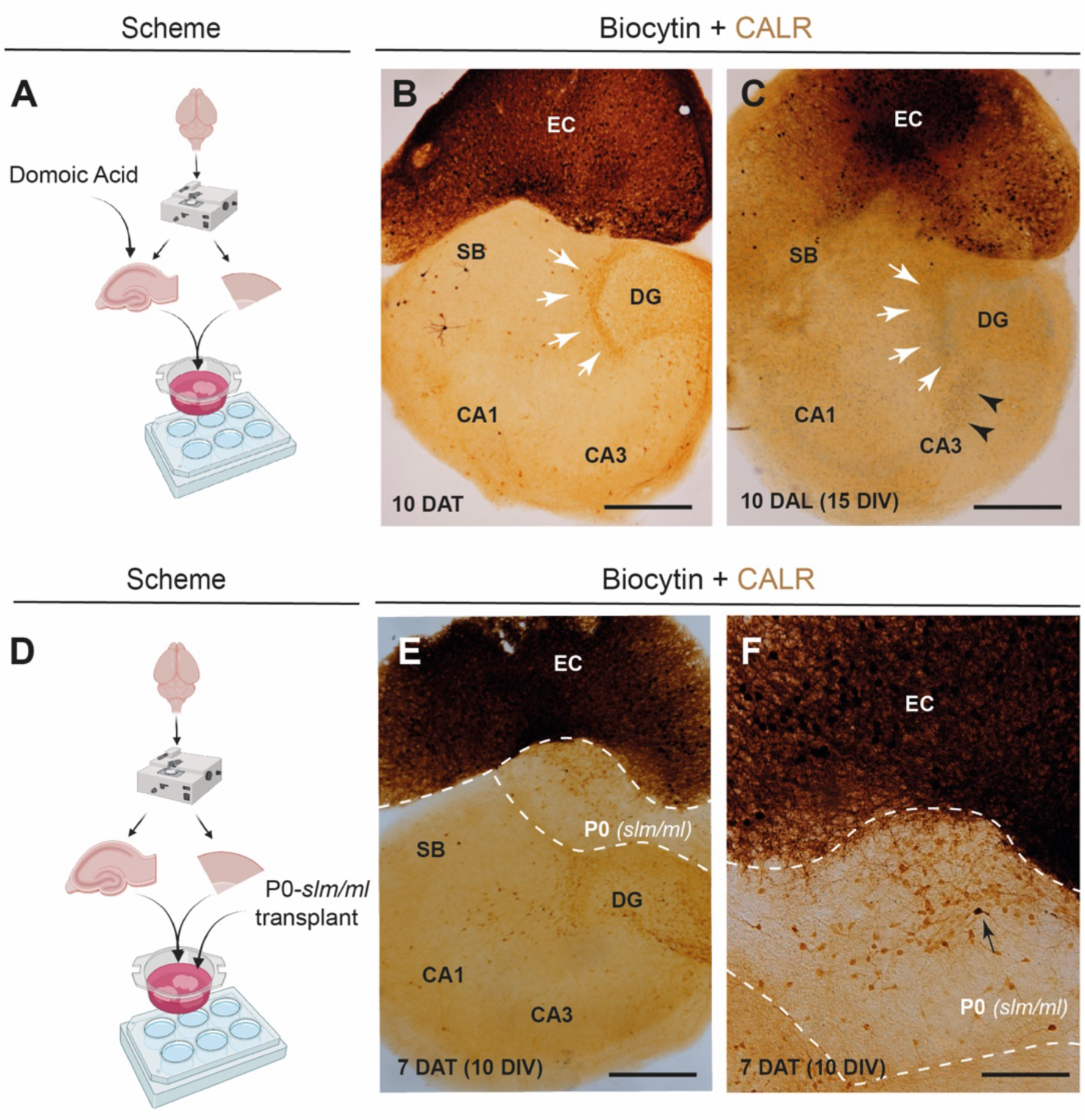
*In vitro* manipulation of the EHC reveals EC intrinsic factors that negatively influenced axon regeneration in aged OECs. A) Scheme illustrating the treatment of the OECs with domoic acid (see Material and Methods for details). **B)** Photomicrograph of a vibratome section of an OEC treated with domoic acid at 1 DIV after 10 DAT. Notice both the absence of EC axons in the hippocampus and the absence of most Cajal-Retzius cells (arrows) after this early domoic acid treatment of the hippocampal slice. **C)** Example of an OEC of the EHC at 10 DAL of the EC axons at 15 DIV. As also illustrated in Figure 1, EC axons are unable to innervate the hippocampus (arrows). In addition, numerous proliferating cells can be observed in the deafferented region of the CA1-3 (arrowheads). **D)** Scheme illustrating the transplantation procedure after EC axotomy. **E-F)** Low **E** and high-power **F** photomicrograph illustrating vibratome sections of two examples of transplanted EHCs ((7 DAT) after EHP axotomy at 10 DIV) with *slm/ml* from P0 hippocampus after Biocytin tracing and Calretinin immunostaining 7 DAT at 10 DIV. The dashed line indicates the axotomy trajectory and delineated the transplanted region. Notice the survival of the transplanted Cajal-Retzius cells in the transplanted *slm/ml*, the presence of residual Cajal-Retzius cells in the lesioned hippocampus and the presence of a retrogradely labeled Cajal-Retzius cell (arrows). In addition, notice the absence of EC axons in the transplanted target and the hippocampal slice. Scale bars: B, C and E: 200 μm; F: 100 μm. Abbreviations as in Figure 1.

### Development of an optogenetic stimulation setup of the EC neurons in EHCs

Cell specific neural stimulation has been used in several paradigms. In our laboratory this stimulation has been developed in several culture lab-on-chip platforms ([72, 77, 88]. More relevantly, our data in 2D dorsal root ganglia (DRG) cultures point that neuronal or chemogenetic stimulation of lesioned DRGs only induce axon regeneration when neurons are cultured on permissive in contrast to inhibitory (containing chondroitin sulfate proteoglycans, CSPG) substrates. CSPG, among other inhibitory molecules, is largely present after EHC axotomy in OECs [59]. Thus, we aimed to explore whether the specific activation of EC neurons might modify both the natural development of the EHC and their regeneration after axotomy. In an elegant study, Hildebrandt-Einfeldt and coworkers demonstrated that optogenetic activation of entorhinal neurons expressing ChR2 was able to induce membrane depolarization in hippocampal granule cells [12]. However, in order to develop these experiments, we wanted first to increase the amount of ChR2-expressing EC neurons in our co-cultures (Figure 4A) and to demonstrate that the optogenetic activation of the EC takes place in our OECs. Thus, we established our AVV infection protocol and developed a specific setup in our Olympus BX61 microscope to demonstrate the effectiveness of the optical treatment (Supplementary Figure 1). In the AVV protocol, the EC slices were double infected overnight the same day of dissection with two different AAV9s encoding ChR2-eGFP and JRCaMP1b (see Material and Methods for details) both genes under the Synapsin promoter (Figure 4B-E). Analysis of the colocalization between eGFP and JRCAMP1b at 10 DIV using JACoP ImageJ plugin [79] determined a Pearsońs coefficient of 0.8348 ± 0.0291 (Mean ± SD) with and Overlap coefficient of 0.9908 ± 0.0095 (Mean ± SD) indicating high level of co-expression (Figure 4E). For stimulaton, a diode laser of 470 nm was placed perpendicular to the optical plane to illuminate the OECs. In addition, the activity measured as Ca^2+^ changes were monitored used the appropriate filter sets that blocked light wavelength lower that 618 nm (Supplementary Figure 1A). In these conditions, we developed short videos of ∼80 sec to determine whether the lateral illumination with the diode laser induced an increase in the intracellular Ca^2+^ of JRCaMP1b- and ChR2 co-infected ECs. In control experiments with JRCaMP1b expression in absence of ChR2, no fluorescence changes were observed after the illumination at 470 nm (Supplementary Figure 1B-D). In Figure 4F we showed one frame of one example of the jRCaMP1b analyzed videos. Due that the observation implies the immersion of our objective in the media was used the putative non-rigid movements in our sample during recording was corrected by using EzCalcium^TM^ and the ROIs were selected by CNMF-E algorithms of the software (Figure 4G). In addition, video time lapses were processed by using Aydin^TM^ to reduce the noise of the recording. After this, selected ROIs after NeuroSeg analysis were analyzed (Figure 4H), and ι1F/F0 of each ROIs was determined (Figure 4I). As observed, after the illumination between 7 and 40 sec (in this example) a transient increase in the Ca^2+^ on jRCaMP1b EC-positive cells (mainly in the border of the slice) was observed due to the location of the 470 led beam (Figure 4I, Supplementary movie 1). Thus, with these experiments we demonstrated that our optical treatment treatment could activate EC neurons expressing ChR2.

**Figure 4.**
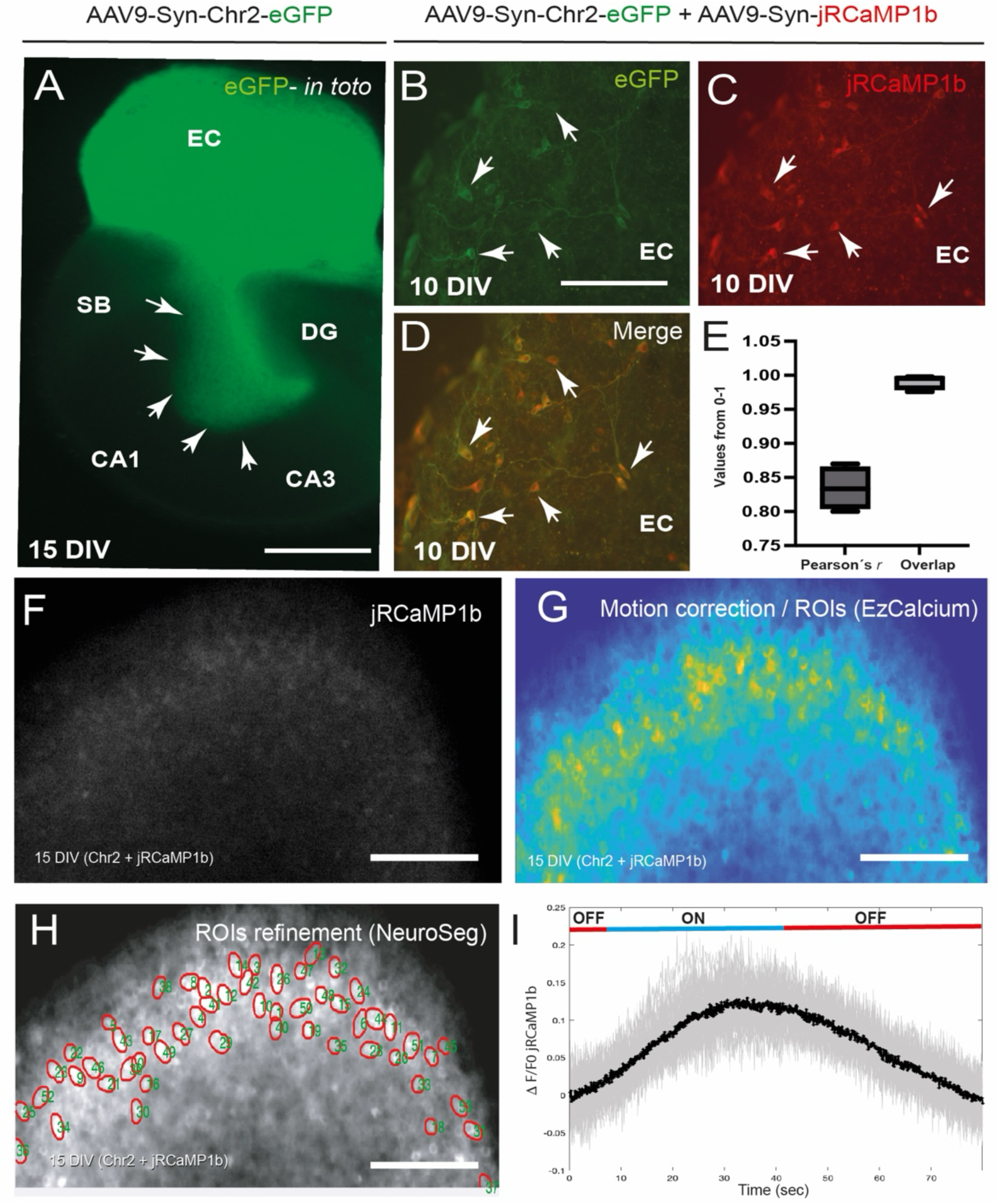
Optogenetic stimulation of developing EHC in OECs. **A)** Whole-mount fluorescence photomicrograph of an OEC infected in the EC with AAV9 expressing ChR2- eGFP. Note that the expression of ChR2 does not impair the proper development and target- specific growth of the EHP (arrows), and only EC axons are fluorescent. **B–D)** High-power photomicrographs of a doubly infected EC with AAV expressing ChR2- eGFP (**B**) and jRCaMP1b (**C**). **E)** Graph illustrating the mean Pearson’s *r* and Overlap values obtained from the analysis of several sections from 8 double-infected cocultures (values are Mean ± SD). **F)** Photomicrographs illustrating a frame from the analyzed videos in the red channel of jRCaMP1b.**G)** Image showing the location of the ROIs detected by EzCalcium using a physics-based lookup table. **H)** Delineation of the selected ROIs after NeuroSeg analysis. **I)** Graph illustrating in gray the changes in *ΔF/F0* calculated for each ROI during the study period. The average of the observed changes is shown in black. At the top, the fraction of time in which the culture was exposed to light (ON) or not exposed (OFF) is indicated. **Scale bars:** A: 200 μm, B–D: 100 μm, F–H: 150 μm. Abbreviations as in Figure 1.

### Optogenetic stimulation of developing EHP largely alters their development in OEC

Considering that, our stimulation protocol was able to enhance EC neuron activity, we aimed first to determine whether this increased activity could change the laminar specific innervation of EC into the hippocampus during its development (Figure 5). In a first set of experiment, we determined that, with the AAV infection protocol, only EC projecting neurons expressing ChR2 were stimulated. Thus, whole mount imaging demonstrates that only treated EC axons innervating the hippocampus express ChR2 during OECs development an example at 15 DIV (see Figure 4). Next, we stimulated a group of EHC infected with AAV expressing ChR2 in the EC during the first 7 DIV (every 12 h). In non-stimulated OECs the development and the layer specific development of the EHP is maintained at 7 DIV (Figure 5A). However, in stimulated cultures we observed that although EC axons were able to innervate the hippocampus, a large number of EC axons showed misrouted trajectories innervating regions other than the *slm/ml* (Figure 5B-E). In the Figure, some examples of Biocytin labelled EHC can be seen. This result was observed in ∼80% of the stimulated co-cultures. This indicate that the stimulation protocol induces an aberrant grow of projecting EC neurons in our OECs.

**Figure 5.**
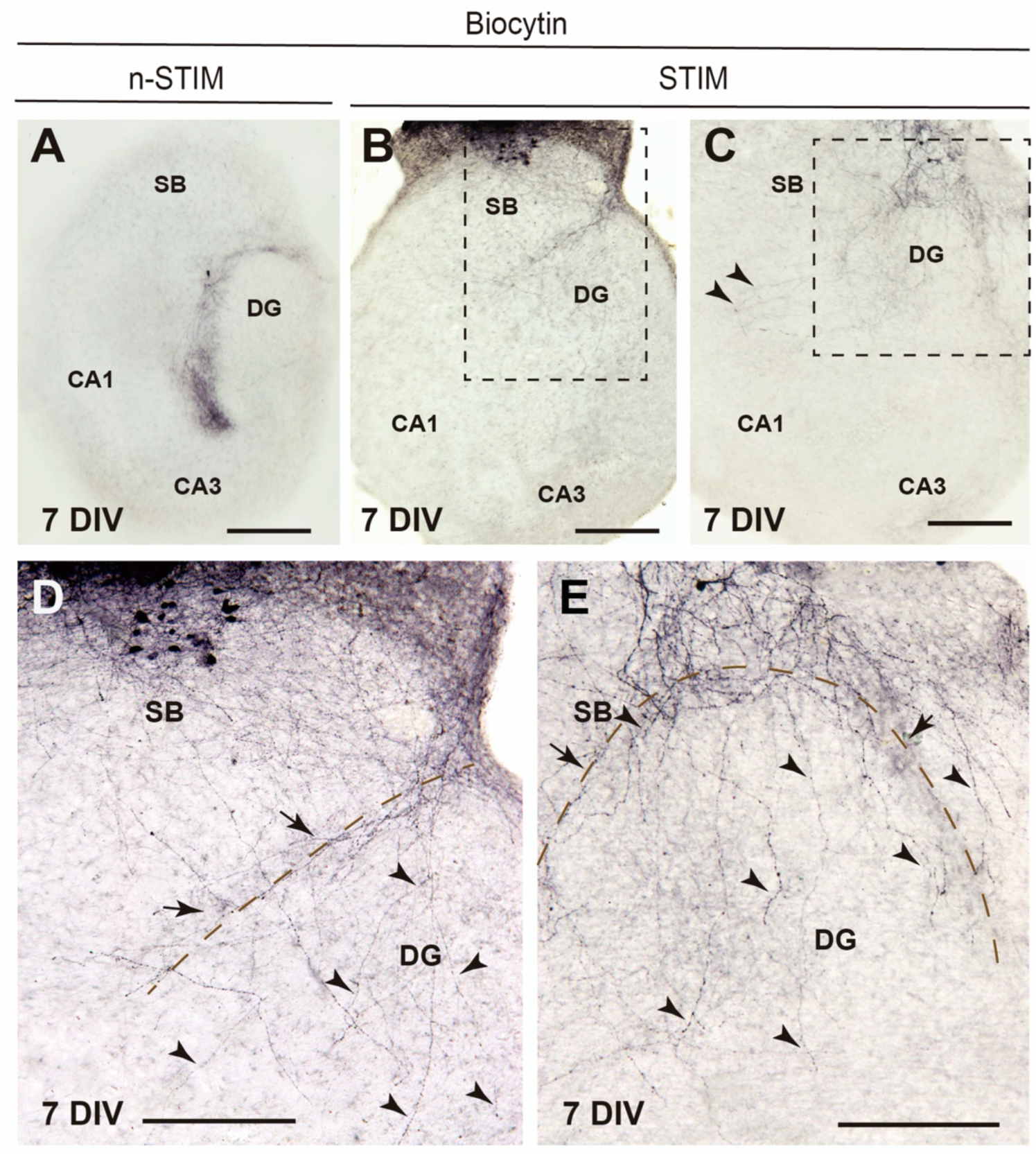
The optogenetic stimulation of developing EHC in OECs impairs target derived signals on growing EC axons leading to misrouted axon innervation. **A)** Vibratome section of a non-stimulated EHC illustrating the “natural” development of the EHP *in vitro.* Please compare this picture with Figure 1B. **B-C)** Examples Biocytin traced EHC at 7DV after the stimulation of EC neurons (se Material and Methods for details). Numerous axons reached the hippocampus showing aberrant trajectories. The dashed Box in **B** and **C** is magnified in D-E respectively. **D-E)** At high magnification, EC-labeled axons (arrowheads in **D** and **E**) reached the hippocampal fissure (dashed line) crossing the subiculum but crossed granule cell layer reaching the hilus and the CA1 and CA3 regions. Scale bars: A-C: 200 μm; D-E: 100 μm. Abbreviations as in Figure 1.

### Optogenetic stimulation of axotomized EHP do not enhance axon regeneration in aged EHC

In a final set of experiments, we aimed to explore whether the stimulation of lesioned EC neurons can enhance their regenerative properties. We performed the axotomy of several OECs at 14 DIV and developed the optical stimulation protocol using our optical platform (Figure 6A-D). Unfortunately, fluorescent observation (Figure 6C) and Biocytin tracing (Figure 6D) renders similar negative results with no EC axons innervating the hippocampal slice. In parallel experiments we analyzed whether the stimulation might affect the survival of projection neurons, and the establishment of the glial scar observed in Figure 2. Results also indicate that the stimulation process do not affect neither neuronal survival nor the generation of the glial scar which might impair the EC axon regeneration (not shown).

**Figure 6.**
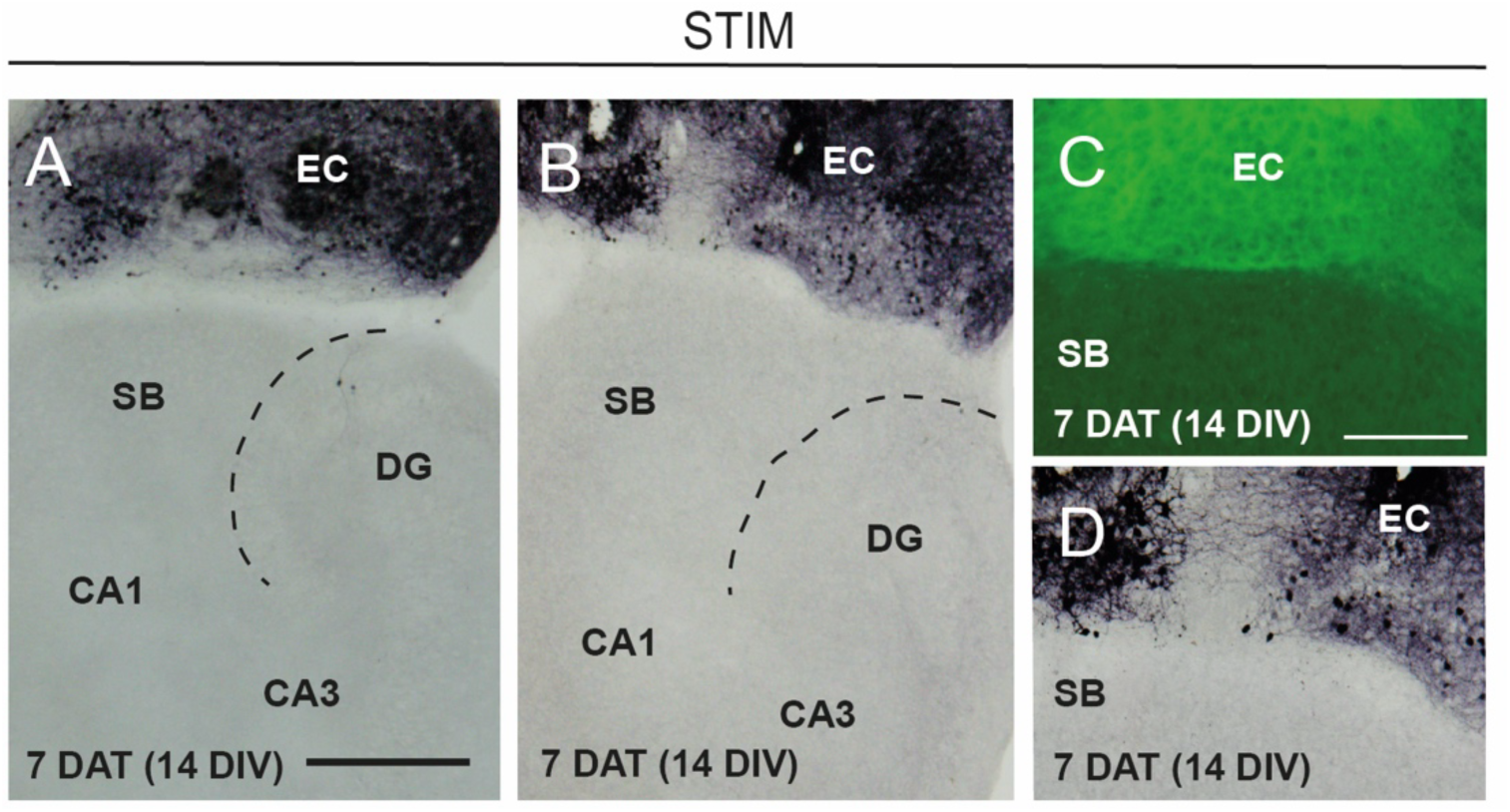
Failure of the optogenetic stimulation of axotomized EHC in promoting axon regeneration in OECs. A-B) Medium (**C-D**) and high magnification photomicrographs illustrating the absence of EC axonal regeneration after the stimulation procedure, analyzed by Biocytin (**A-B** and **D**) or eGFP fluorescence **(C)**. Scale Bars: A: 200 μm pertains to B; C: 100 μm pertains to D. Abbreviations as in Figure 1.

### Decreased expression of axonal growth-related genes under enhanced activity of the EC neurons

Due that our experiments point to a dysregulation of the pathfinding properties of developing EC axons and the absence of EHP regeneration after lesion. We aimed to explore whether the stimulation protocol of the EC neurons changes particular genes expression involved in axonal growth. Thus, levels of *GAP43*, *CAP23, BDNF, NGF* and *SP11RA*, were analyzed in axotomized EHC after stimulation and its expression was compared to non-stimulated cultures using *GADPH* as housekeeping control gene. Although the number of processed cultures were low (n = 4 at each group), results indicate some tendencies in down regulating gene expression between stimulated *vs* non-stimulated EC neurons (Figure 7A). These results reinforce the notion that although we cannot completely rule out that glial scar-derived inhibitory molecules might be involved, the stimulation protocol developed *in vitro* is not robust enough to induce positive changes in gene expression to enhance are axon regeneration in EHC after axotomy. In fact, our data points that the forced stimulation of EC activity is detrimental for regeneration and, to the correct development of the EHC *in vitro*.

**Figure 7.**
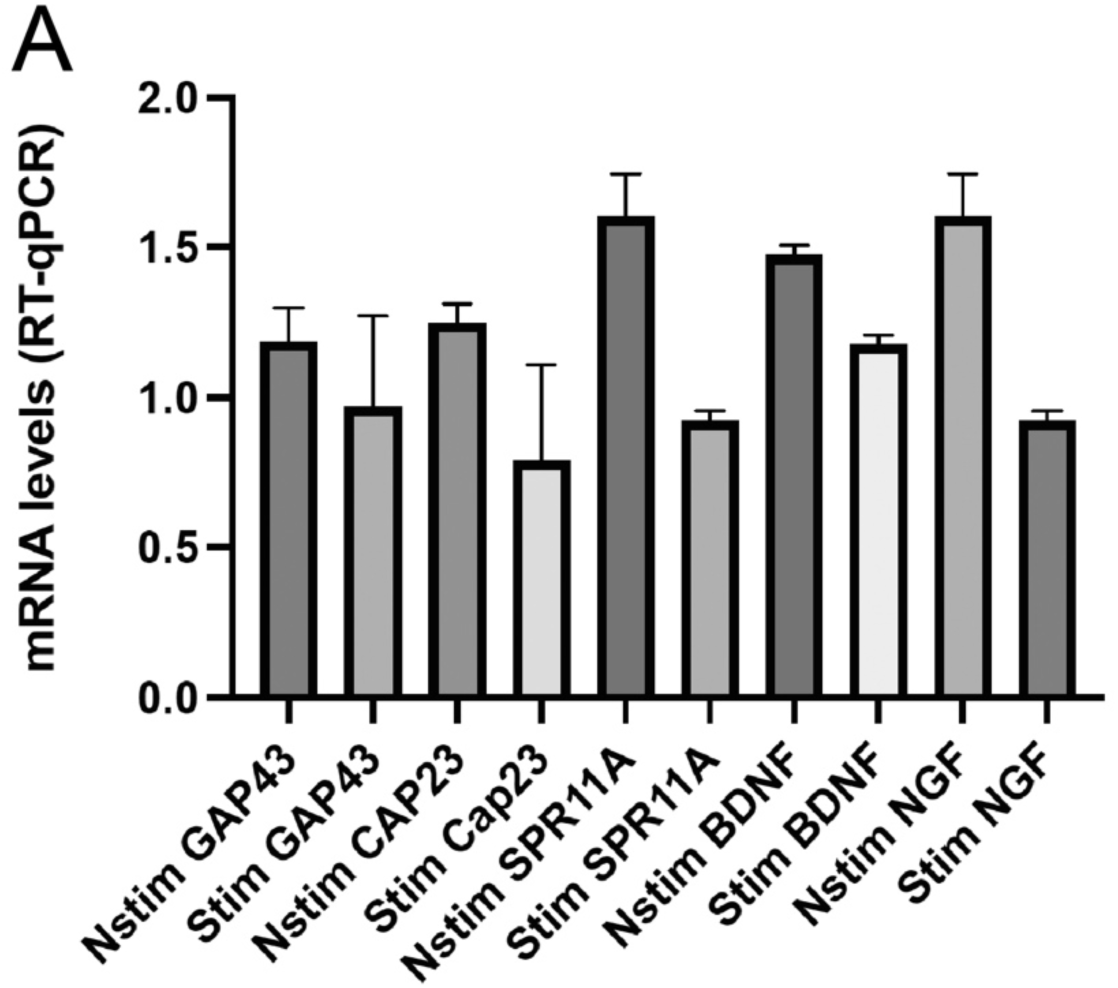
Changes in genes expression in the axotomized EC after light stimulation. Gene expression values analyzed by RT-qPCR (see Materials and Methods for details). Although the number of cases is limited, there is a trend toward reduced expression of the analyzed genes following stimulation. The observed values have been normalized to *GAPDH* expression levels.

## Discussion

Axonal wiring in mammals is controlled by genetic cascades as well as by intrinsic activity-dependent development and refinement of connections [89]. These activity dependent development has been described, among other, in the optic tectum [90], motor neurons [91, 92], midline callosal axons [93] or mouse somatosensory and visual cortices [94, 95]. From several years, it has been considered that spontaneous activity has been shown to be important in refining neural projections once axons have reached their targets, but early pathfinding events have been thought to be activity independent. In other words, the general consensus in the field has been that early events such as axon growth to the target depends on a complex series of molecular signals but is independent of neural activity [96, 97]. It is currently known that this basal activity in projecting neurons plays a fundamental role in the proper recognition of target regions and their subsequent innervation and maturation (e.g., [98, 99]). Thus, a physiological balance must be stablished to modulate both axonal growth and target recognition. In a pioneer study during spinal cord development, over-stimulated motor axons correctly executed the binary dorsal-ventral pathfinding decision but failed to make the subsequent pool-specific decision to target to appropriate muscles [92]. In addition, decreasing the frequency of spontaneous bursting episodes results in dorsal-ventral pathfinding errors for motor projections (see [100] for review). The EHP is one of the best studied pathfinding models and is fundamental to the establishment of archicortex connectivity in the nervous system. However, it is not known whether EHP development requires intrinsic activity in axons, or what regulates that activity. Furthermore, a mechanism linking neuronal activity and gene expression has not been identified in relation to axon pathfinding within the EHP. In our experiments, an increase in endogenous spontaneous activity in developing EC neurons impairs their ability to recognize targets in the developing hippocampus. This effect is associated with EC-projecting neurons nor the target cells, due to their exclusive expression of ChR2. This is relevant, as it supports published studies indicating that changes in neuronal activity can affect axonal pathfinding in the visual and motor systems (see [101–104] for additional references).

Although more experiment needs to be developed, in our experiments we observe that several genes involved in axon grow and regeneration (e.g., *GAP43*) among others was decreased in stimulated axotomized cultures. Although the role of GAP43 in developing axons has been largely studied (see [105] for review), a recent study indicates that the decrease of GAP43 in hippocampus impairs target specificity and function of mossy fibers [106]. Similarly, some years ago, the role of the neural activity as consolidator of hippocampal inhibitory circuit in the hippocampus was described [107]. Thus, neuronal activity regulates the density of inhibitory synapses made by postnatal hippocampal interneurons, and BDNF could mediate part of this regulation. This regulation of the density of inhibitory synapses could represent a feedback mechanism aimed at maintaining an appropriate level of activity in the developing hippocampal networks, including the EHP [107]. Similar defects during cortical development and axonal wiring has been described for deficits in BDNF [108, 109] or NGF [110]. In fact, some regenerative associated genes are dependent on neurotrophin expression (e.g., [111] see [112] for review). This is also of relevance since some of these neurotrophins (e.g., NGF) can act on growing or regenerating neurons (e.g., [113]) but also can be released anterogradely in order to enhance correct target recognition and synaptic function [114].

Taking this into account, we can consider that the gene expression decrease induced by the stimulation protocol largely impair the pathfinding and the target recognition of EC axons during regeneration and most probably during development. We consider that this is of relevance since our previous results [72], also point in this direction. Additionally, and from a regenerative strategy point of view, sprouting of spared axons, instead of injured ones, is also responsible for recovery after increasing neuronal activity by exercise and/or electrical stimulation (see [115] for review). In agreement to that, success of these approaches is only significant in incomplete injuries with increased number of spared axons. Incomplete injuries might also help explaining why in our model (complete lesion of the EHP) we did not observe recovery after optogenetic stimulation while others did on lesioned spinal cord (e.g., [116]) as different injuries (incomplete lesion) were used. Moreover, complete injuries also present larger glial scars (as observed in EHC *in vitro*), and therefore greater accumulation of growth inhibitory molecules [59, 73, 74]. On the other hand, the expression of the receptors for most of these molecules are maintained in the project EC neurons after axotomy [117, 118], increasing their negative effects that the stimulation protocol cannot overcome. In parallel, our stimulation strategy does not affect the generation of the glial scar in axotomized EHP. This raised the question of whether modulating both glial scar formation as demonstrated by [119] and neuronal activity triggering the axotomized projecting neuron from an apparent resting stage (e.g., as Purkinje cells [120] to a pro-regenerative stage will render better results if take into account that EC neurons do not disappeared after axotomy *in vitro* and *in vivo* [69, 121].

## Acknowledgements

We would like to thank all the members of the laboratory for their valuable comments on the study. We would also like to thank Renato Eduardo Yanac Huertas and Oscar Castaño Linares (UB) for the help in developing *ad hoc* Python 3.9 based graphic user interphase (GUI) for calcium analysis (Analyca) and our collaborators in providing Matlab^TM^ algorithms and reagents for this study. We also wish to thank the members of the Electron Microscopy Service at the University of Barcelona and the staff of Core Facilities and the MicroFabSpace (IBEC) for the technical support provided.

