## Supplementary material for "Optogenetic activation of entorhinal projection neurons alters the target recognition and circuit development without enhancing axon regeneration after axotomy in organotypic slices": Suppl. Figure 1

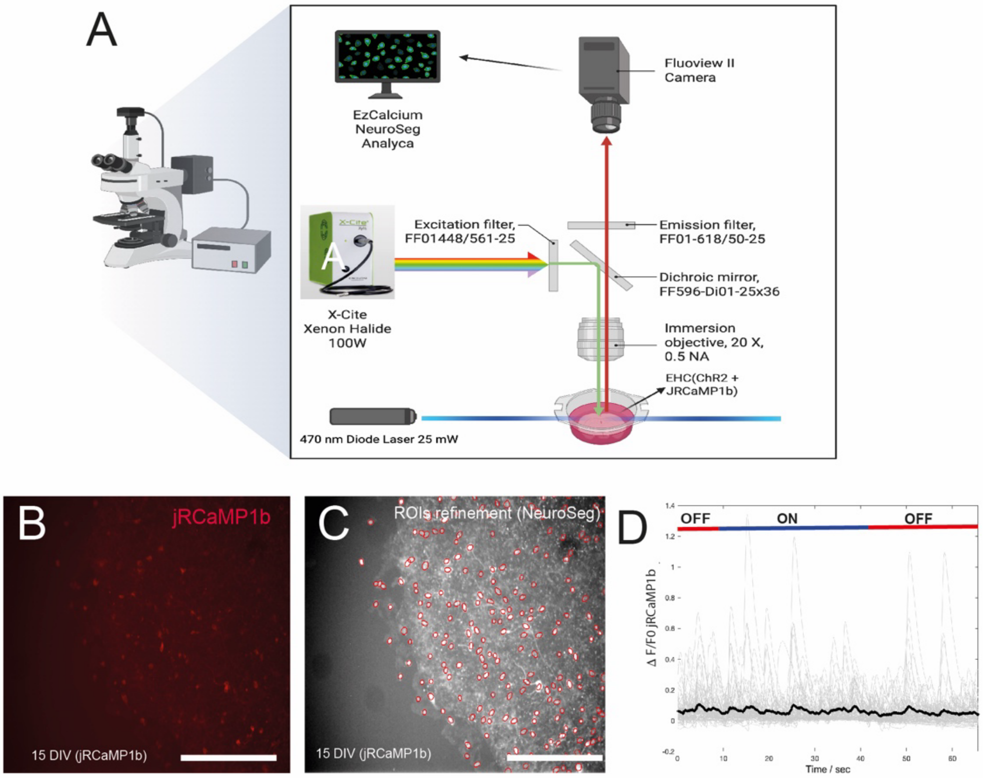


SUPPL. FIGURE 1

**Supplementary Figure 1.**

**Experimental setup used to study neuronal activity after stimulation.** **A)** This diagram details the filters used in the study. As shown, the light source is a 100W X-Cite lamp. After passing through the optical filters, only wavelengths above 591 nm are allowed through. This light excites jRCaMP1b, and the fluorescence emitted by the GECI passes through a selective filter that allows wavelengths above 618 nm. For ChR2 stimulation, a pulsed 470 nm laser diode is placed at the side of the transwell to avoid aligning with the direction of the light generated by the X-Cite system. This laser can be switched on and off manually by the researcher. The resulting fluorescence is captured by the Fluoview II camera and later analyzed using the software described in the manuscript.**B)** Example of an EC infected with jRCaMP1b but not with ChR2. **C)** After analysis with EzCalcium and NeuroSeg, the ROIs selected in this sample are shown. **D)** As indicated in Figure 4, the graph displays the changes in $\Delta F/F0$ for the different ROIs analyzed, with the average shown in black over time. As can be seen, in the absence of ChR2, no significant changes in $\Delta F/F0$ values are observed.
